# A genetic variant alters the secondary structure of the lncRNA H19 and is associated with Dilated Cardiomyopathy

**DOI:** 10.1101/2021.01.22.427739

**Authors:** Leonie Martens, Frank Rühle, Anika Witten, Benjamin Meder, Hugo A. Katus, Eloisa Arbustini, Gerd Hasenfuß, Moritz F. Sinner, Stefan Kääb, Sabine Pankuweit, Christiane Angermann, Erich Bornberg-Bauer, Monika Stoll

## Abstract

lncRNAs are at the core of many regulatory processes and have also been recognized to be involved in various complex diseases. They affect gene regulation through direct interactions with RNA, DNA or proteins. Accordingly, lncRNAs structure is likely to be essential for their regulatory function. Point mutations, which manifest as SNPs (single nucleotide polymorphisms) in genome screens, can substantially alter their function and, subsequently, the expression of their down-stream regulated genes. To test the effect of SNPs on structure, we investigated lncRNAs associated with dilated cardiomyopathy. Among 322 human candidate lncRNAs we demonstrate first the significant association of a SNP located in lncRNA H19 using data from 1084 diseased and 751 control patients. H19 is generally highly expressed in the heart, with a complex expression pattern during heart development. Next, we used MFE (minimum free energy) folding to demonstrate a significant refolding in the secondary structure of this 861 nt long lncRNA. Since MFE folding may overlook the importance of sub-optimal structures, we showed that this refolding also manifests in the overall Boltzmann structure ensemble. There, the composition of structures is tremendously affected in their thermodynamic probabilities through the genetic variant. Finally, we confirmed these results experimentally, using SHAPE-Seq, corroborating that SNPs affecting such structures may explain hidden genetic variance not accounted for through genome wide association studies. Our results suggest that structural changes in lncRNAs, and lncRNA H19 in particular, affect regulatory processes and represent optimal targets for further in-depth studies probing their molecular interactions.

## 1 Introduction

LncRNAs are defined as RNA molecules which are > 200 nucleotides (nt) long and, usually, not translated into proteins [1]. LncRNAs have garnered much attention over the last decade since comparative eukaryotic genomics and transcriptome analyses demonstrated that lncRNAs are abundant and often play important roles for gene regulation in general [2, 3, 4]. So far, lncRNAs have been implicated in several biological and developmental functions through regulation of gene transcription, post-transcriptional RNA processing and chromatin modification [5, 6, 7, 8, 9]. Naturally, several lncRNAs have also been linked to various pathological processes [10, 11].

In the context of comparative human genomics, genome-wide association studies (GWAS) have provided a wealth of genetic associations for common complex diseases. Surprisingly, around one third of all SNPs in the most recent version of the GWAS catalog [12] are located in annotated lncRNAs. Accordingly, lncRNAs are emerging as prime candidates for explaining the missing heritability of complex traits. Several prominent lncRNAs have been described in the context of the cardiovascular system and its diseases [13, 14, 15, 16, 17] which account for the highest mortality in the developed world, exceeding by far the annual deaths from cancer [18, 19]. However, in most cases, the functional and structural description of implicated lncRNAs is rudimentary at best and suitable functional assays are commonly lacking. Since lncRNAs do not code for a protein, their structure is assumed to be essential for their function. Certain RNAs can undergo strong structural changes upon single nucleotide substitutions [20, 21, 22, 23] and it is likely that a group of SNPs affect lncRNA function accordingly. Indeed, a recent study investigating the transcriptome of a family trio (father, mother, child)[22] reported that 15% of the transcribed SNPs altered RNA secondary structure. Furthermore, structure-altering SNPs were linked to altered gene expression and a set of disease phenotypes. These SNPs are designated as RiboSNitches [23]. So far, there are only few well characterized functional examples of structure altering SNPs in regulatory RNAs. One such example was described for a RiboSNitch which, in the wake of a SNP, can no longer attach to a regulatory protein (IREBP, Iron Response Element Binding Protein). As a consequence, the FTL gene, which encodes a subunit of the Ferritin complex, can no longer store excess iron which leads to hyperferritinemia cataract syndrome [24, 25], causing clouded eye lenses at early ages.

Unfortunately, predicting and determining RNA structures remains notoriously difficult and almost impossible for long molecules such as lncRNAs. However, a recent surge in experimental techniques coupled with next generation sequencing (NGS) has significantly improved predictions. These techniques comprise enzymatic and chemical approaches, depending on the reagent used to modify certain nucleotides before probing the structure. One prominent chemical approach is called SHAPE (selective 2-hydroxyl acylation by primer extension)-Seq [26], which binds to unpaired nucleotides and interrupts the reverse transcription process. Subsequent sequencing and analysis of the differences in read length compared to a control is used for computational structure prediction, significantly increasing the reliability [27].

Here, we set out to investigate the influence of dilated cardiomyopathy (DCM) associated RiboSNitches on lncRNA structure and function. First, we identified 14 disease associated SNPs located in several lncRNA transcripts. Next, we investigated them for their structure through MFE structure prediction algorithms. To further support these results, we also compared the change in the suboptimal structure ensemble. Here, the lncRNAs Carmn, H19 and MLIP-AS1 demonstrated that the according SNP had a massive effect on the relative probabilities of likely sub-optimal structures. Finally, we used SHAPE-Seq analyses to experimentally confirm the *in silico* predictions of these potential RiboSNitches.

Indeed, we found the SNP rs217727, located in the lncRNA H19, to be disease associated and inducing a remarkable shift in the structural ensemble. This may impair lncRNA function, and thus contribute to the pathogenesis of complex diseases such as heart failure.

## 2 Materials and Methods

### 2.1 Association Analysis

The samples for sequencing and genotyping were part of the German National Genome Research Network (NGFN) call “The genetics of heart failure - from populations to disease mechanisms”. First, 96 DCM samples were screened for genetic variants in lncRNAs. Customized probes for the Illumina Nextera rapid capture enrichment protocol were used for the lncRNA exonic regions. Reads were mapped to the hg19 build using BWA mem [28], followed by variant calling using GATK 3.7 [29]. Association analysis was then performed using Plink 1.7 [30] logistic regression assuming an additive model with sex as covariate and matching controls from the 1000 Genomes project[31]. Promising variants were subsequently confirmed by genotyping of a larger cohort comprising 1084 DCM cases and 751 disease-free controls. The study was conducted in accordance with the principles of the Declaration of Helsinki. All participants of the study gave written informed consent and the study was approved by the local ethic committees at the participating study centers.

### 2.2 Secondary Structure Predictions

RNA secondary structure prediction algorithms are based on thermodynamics and most often the minimum free energy (MFE) structure is determined. The MFE calculation is aiming to determine the predicted structure with the lowest free energy, since it is assumed that the lower the value, the more stable and likely the structure. For this, most algorithms are using the dynamic programming approach based on Zuker et al. [32]. Here, MFE structure predictions were performed using the ViennaRNA package RNAfold [33]. Since the MFE structure is not necessarily the only naturally occurring structure, we also considered suboptimal structures. For this, structures from the Boltzmann distribution were sampled with the ViennaRNA package RNAsubopt [33]. The resulting structures were then transformed into a binary vector, followed by t-SNE for visualization (according to [25, 34]).

### 2.3 In vitro SHAPE-Seq

Genetic variants that were significantly associated with our phenotype and were predicted to influence the structural properties of a lncRNA were subjected to validation through SHAPE-Seq structural probing. First, the RNA transcripts were generated in vitro from a ThermoFisher Gene Synthesis Plasmid with the T7 RNA polymerase. In vitro SHAPE-Seq was performed according to the protocol by Watters et al. [26] using the 1M7 reagent. The samples were sequenced 80 cycles in a paired-end mode with the Next.Seq500 system and v2 chemistry. The resulting reads were analyzed with the accompanying Spats software for generating a reactivity profile. The resulting reactivities were then incorporated as an additional constraint during secondary structure prediction. RNAStructure [35] uses the approach first suggested by Deigan et al. [36] which transforms the reactivity information into pseudo-free energy using the equation ΔG = *m ln*[*r*+1]+*b*, where *m* and *b* are heuristically determined comparing probing data to crystallization structures, and *r* are the reactivity values. The SHAPE-probing information guided sampling of the structural ensemble was performed through the RNAStructure [35] pipeline consisting of Rsample, stochastic, and RsampleCluster. This pipeline is based on the algorithm by Ding and Lawrence [34, 37] for sampling and centroid determination and diana [38] followed by Calinski-Harabasz index [39] for finding the optimal *k* for clustering. In order to visualize the structures, the t-SNE [40] algorithm was utilized for dimensional reduction through the R package Rtsne [41],

## 3 Results

We found 14 associated SNPs (p ≤ 0.05, coverage ≥ 30) that overlapped with lncRNA transcripts (limited to 1000 nt length). We prioritized transcripts that were expressed in human myocardium samples (SRR1957191, SRR3151752, SRR3151758) [42, 43] and demonstrated a larger change in the structural ensemble. Genotyping of selected RiboSNitches (rs13158382 (Carmn), rs217727 (H19) and rs12527071 (MLIP-AS1)) in our confirmation case-control cohort, confirmed the association between DCM and rs217727, located in lncRNA H19.

### 3.1 Structural prediction

The MFE structure prediction of the H19 reference and rs217727 RiboSNitch sequences showed a local change in the structure located around the altered nucleotide (Figure 1).

**Figure 1:**
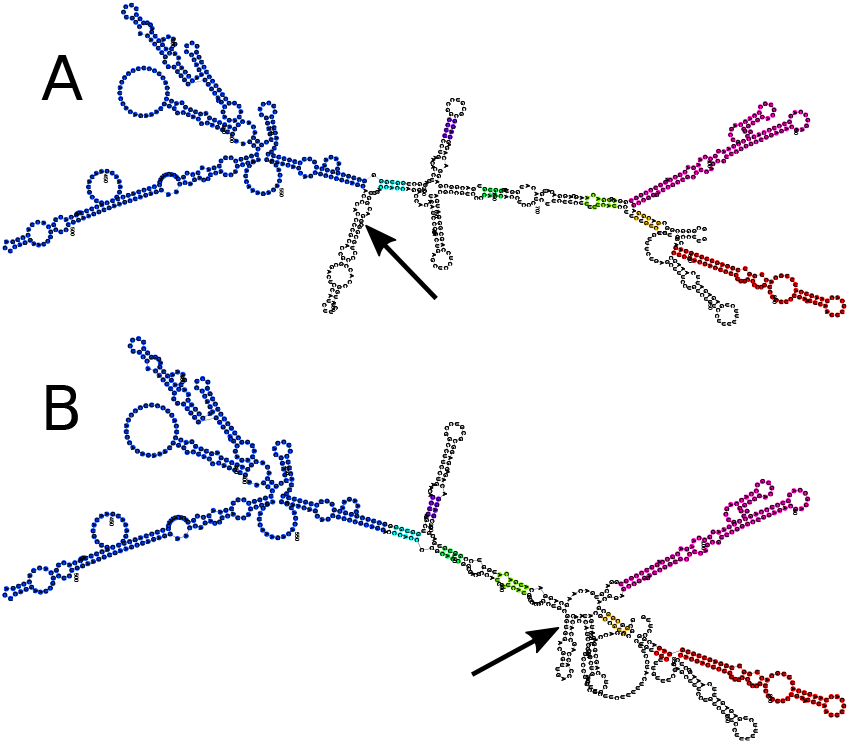
Minimum Free Energy (MFE) structure of the lncRNA H19. Secondary structure prediction was done using RNAfold. **A** Is the reference structure and B is the structure changed through the potential RiboSNitch rs217727. Arrows indicate the changed nucleotides and identical colors highlight unaffected substructures. Reference structure free energy: −306.50 kcal/mol; RiboSNitch structure free energy: −308.50 kcal/mol.

Because most RNAs can assume multiple or different structures from the MFE structure itself, these predictions are not completely accurate particularly for long RNA structures. Therefore, we considered the ensemble of sub-optimal RNA structures. The entire ensemble of structures, sampled from the Boltzmann distribution, shows a clear shift, which is another indicator of a potential RiboSNitch (Figure 2).

**Figure 2:**
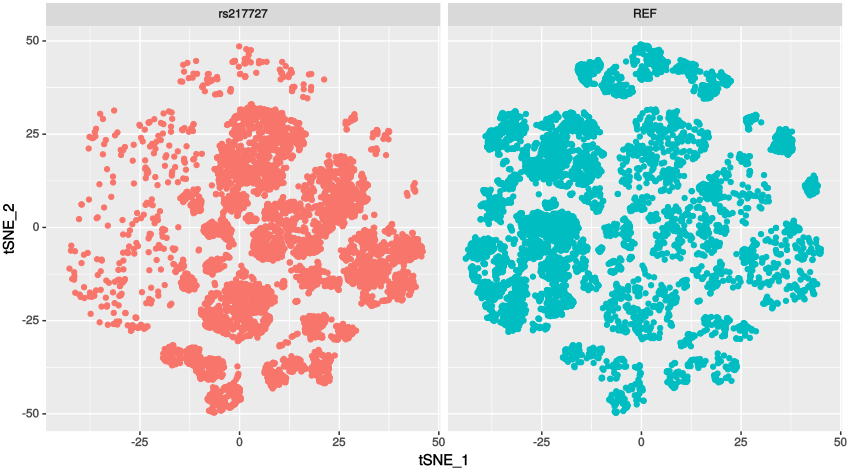
tSNE of suboptimal structures for H19 sampled from the Boltzmann ensemble (n=10000). **Left** are the structures for the RNA with the genetic variant rs217727 and **Right** of the reference RNA. The shift shows that due to the genetic variant, other structures become more likely, which can lead to a change of function.

### 3.2 Structural probing

Although computational prediction of secondary structures has been widely used, the accuracy is often insufficient [44] and decreases for longer sequences. Therefore, experimentally derived data is needed to guide prediction algorithms. After successfully establishing and following the SHAPE-Seq protocol for H19 with the wildtype and the minor allele of rs217727, we received between 9 and 19 million paired-end reads. This was sufficient for the 861 nucleotides long sequence of lncRNA H19 to reach an average coverage of 5186.54 (minimum coverage of 106.5). These resulting reactivities were then incorporated during structure prediction. When assessing the accuracy of the predicted MFE structure compared to the SHAPE-Seq aided prediction through the RNAstructure [35] scorer function, we found very low accordance (sensitivity 45.95%; positive predictive value (PPV) 41.98%). As a next step, we investigated the structural ensemble, while taking the SHAPE-seq data into account. In Figure 3, suboptimal structures show clear differences between the reference structures and the RiboSNitch structures. Cluster analysis identified two and three clusters, respectively. Each cluster was assigned a centroid structure, which is the most representative structure.

**Figure 3:**
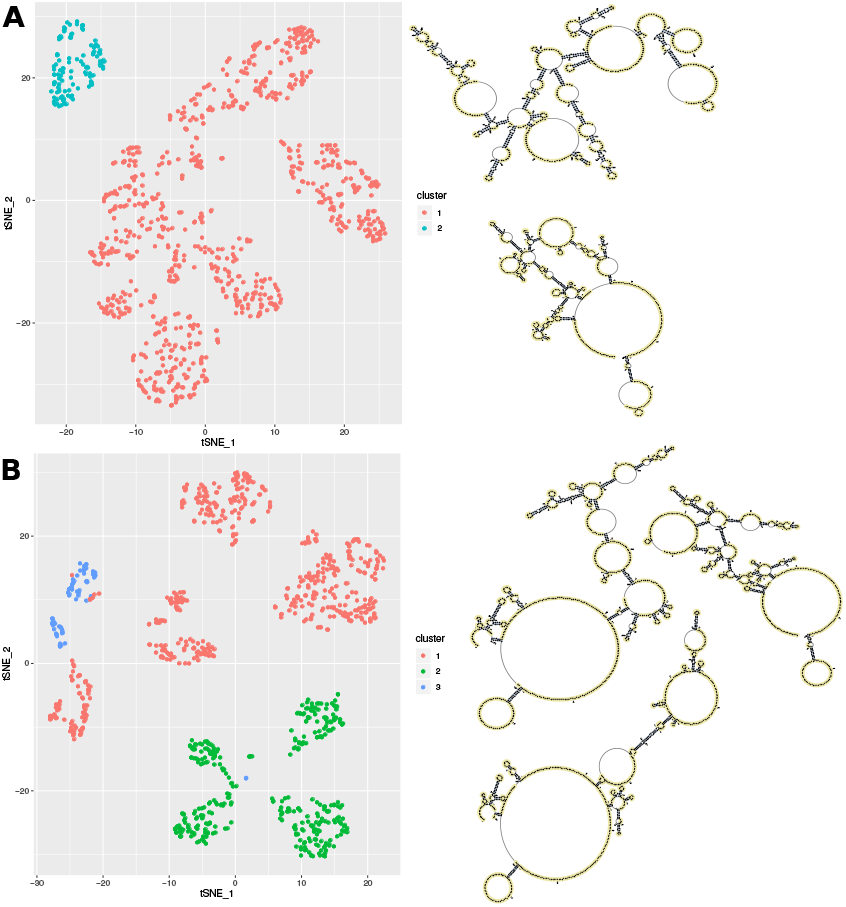
Visualization of the structural ensemble taking the SHAPE-seq data into account. For this 1000 structures were sampled from the Boltzmann distribution, that were colored by their optimal cluster affiliation. t-SNE was used for dimensionality reduction and plotted in 2D. **A** Cluster analysis of the reference structures reveals two distinct clusters with different centroid structures. **B** Cluster analysis of the RiboSNitch structures reveals three distinct clusters with different centroid structures.

## 4 DISCUSSION

Here, we investigated the influence of genetic variants on the abundant group of functional lncRNAs. In particular, we were interested in SNPs affecting secondary structure of lncRNAs, potentially contributing to disease. DCM associated SNPs were investigated with respect to their potential as RiboSNitches. Since computationally predicted structures alone are not sufficiently reliable, we combined sophisticated computational and experimental methods. We found that the variant rs217727, located in the lncRNA H19 was not only significantly associated with the DCM disease phenotype, but also highly affected RNA secondary structures. In a previous study, H19 together with the genetic variant rs217727 was associated with an increased risk of coronary artery disease in a Chinese population [45]. The lncRNA H19 is evolutionary conserved across several species, highly expressed in the adult heart and located in an imprinted gene cluster, which also includes genes like IGF2. Whereas H19 is solely expressed from the maternal allele, the growth factor IGF2 is expressed paternally. Despite their reciprocal expression, both genes are still able to regulate each other through the methylation machinery. For example, a mutation in the CTCF target site upstream of H19 has been shown to lead to loss of IGF2 imprinting [46] and such a loss of imprinting in murine embryos resulted in major cardiac abnormalities [47]. Furthermore, H19 has been found to be dysregulated in cardiac hypertrophy and heart failure [48, 49]. In particular, it was shown that knock-down of H19 induced cardiomyocyte hypertrophy, whereas over-expression reversed the effect [49]. H19 can also function as a microRNA-sponge in important regulatory mechanisms involved in cardiac fibrosis. It is competing through binding of miR-455 with connective tissue growth factor (CTGF) and knockdown of H19 confirmed the antifibrotic influences of miR-455 and CTGF amongst others [50]. Thus, in case of a conformational change of the RNA due to a genetic variant, the ability of acting as a microRNA-sponge would be impaired and the expression of the microRNA would consequently increase. This is in accordance with a previous study investigating the miRNOME by deep sequencing of the human heart, where miR-445 was upregulated in DCM patients compared to healthy controls [51]. Another direct target of H19 in the development of fibrosis was identified by Tao et al. [52]. H19 negatively regulates DUSP5 through epigenetic mechanisms and decreased DUSP5 levels lead to an increase in proliferation of cardiac fibroblasts via the MEK/ERK signaling pathway. Additionally, H19 plays an important role in regulating apoptosis. Experiments in rat models for adriamycin-induced DCM showed that H19 is upregulated, whereas knockdown decreased cardiomyocyte apoptosis, resulting in improved left ventricular structure and function [53]. In contrast, Li et al. [54] reported that overexpression of H19 in diabetic cardiomyopathy improved left ventricular function by reducing oxidative stress, inflammation, and apoptosis. Fibrosis and apoptosis are relevant mechanisms in cardiac remodeling and are important processes with an impact on ventricular function in cardiac pathologic conditions [55]. In fact, genetic variants located in H19 and its host microRNA miR-675 have been shown to play a significant role in cardiac pathophysiology in previous studies, increasing the risk for developing hypertrophic cardiomyopathy [56] and coronary artery disease [45]. Since H19 is non-coding, the only way its variants may influence downstream regulation is through structural changes resulting in altered lncRNA function. For example, in rat models of diabetic cardiomyopathy, RNA-binding protein immunoprecipitation experiments were able to demonstrate the ability of H19 to bind directly to EZH2, which is part of the Polycomb Repressive Complex 2 (PRC2), affecting the epigenetic regulation of DIRAS3, a growth suppressor [57]. The murine and human H19 was shown to be able to act as an anti-hypertrophic lncRNA through interaction with PRC2, demonstrating its potential for gene therapie in cardiac hypertrophy [58]. In conclusion, we identified the secondary structure of H19, a lncRNA of importance for cardiovascular function and disease. Further we demonstrated, that a RiboSNitch, associated with the disease phenotype of DCM, significantly alters this structure. It is likely that through a structural change the essential region becomes inaccessible for binding interactions, potentially contributing to a pathological outcome. Further validation of the genetic variant in vivo demonstrating the altered lncRNA interactions in animal models would be of great interest.

## 5 Data Availability

All relevant data are accessible through BioProject ID PRJNA614999.

## 6 Acknowledgements

We would like to acknowledge the excellent technical assistance by the staff of the Core Facility Genomics of the Medical Faculty at the University Münster. This work was supported by the fund Innovative Medical Research of the University of Münster Medical School to F.R. (RÜ121510) and the Deutsche Forschungsge-meinschaft (DFG, German Research Foundation) - 281125614/GRK 2220 to M.S.. B.M. is supported by an Excellence Fellowship of the Else Kröner Fresenius Foundation.

